# Acoustic contamination of electrophysiological brain signals during speech production and sound perception

**DOI:** 10.1101/722207

**Authors:** Philémon Roussel, Gaël Le Godais, Florent Bocquelet, Marie Palma, Jiang Hongjie, Shaomin Zhang, Philippe Kahane, Stéphan Chabardès, Blaise Yvert

## Abstract

A current challenge of neurotechnologies is the development of speech brain-computer interfaces to restore communication in people unable to speak. To achieve a proof of concept of such system, neural activity of patients implanted for clinical reasons can be recorded while they speak. Using such simultaneously recorded audio and neural data, decoders can be built to predict speech features using features extracted from brain signals. A typical neural feature is the spectral power of field potentials in the high-gamma frequency band (between 70 and 200 Hz), a range that happens to overlap the fundamental frequency of speech. Here, we analyzed human electrocorticographic (ECoG) and intracortical recordings during speech production and perception as well as rat microelectrocorticographic (µ-ECoG) recordings during sound perception. We observed that electrophysiological signals, recorded with different recording setups, often contain spectrotemporal features highly correlated with those of the sound, especially within the high-gamma band. The characteristics of these correlated spectrotemporal features support a contamination of electrophysiological recordings by sound. In a recording showing high contamination, using neural features within the high-gamma frequency band dramatically increased the performance of linear decoding of acoustic speech features, while such improvement was very limited for another recording showing weak contamination. Further analysis and *in vitro* replication suggest that the contamination is caused by a mechanical action of the sound waves onto the cables and connectors along the recording chain, transforming sound vibrations into an undesired electrical noise that contaminates the biopotential measurements. This study does not question the existence of relevant physiological neural information underlying speech production or sound perception in the high-gamma frequency band, but alerts on the fact that care should be taken to evaluate and eliminate any possible acoustic contamination of neural signals in order to investigate the cortical dynamics of these processes.

## 1 Introduction

The development of brain-computer interfaces (BCI) to restore speech (Guenther et al., 2009; Brumberg et al., 2010; Leuthardt et al., 2011) is a long-term quest that seems within possible reach. Several advances have indeed been made over the past decade regarding the decoding of intracranial brain signals underlying either speech perception (Pasley et al., 2012; Chan et al., 2013; Pasley and Knight, 2013; Fontolan et al., 2014; Hyafil et al., 2015; Yildiz et al., 2016; Akbari et al., 2019) or production (Bouchard et al., 2013; Martin et al., 2014, 2016; Mugler et al., 2014; Cheung et al., 2016; Chartier et al., 2018; Anumanchipalli et al., 2019), and most recent works have tackled with noticeable success the prediction of continuous speech from ongoing brain activity. Because of the difficulty to record from individual neurons with microelectrodes inserted in speech areas (Bartels et al., 2008; Kennedy et al., 2011; Tankus et al., 2012; Chan et al., 2013), most of speech decoding studies use field potential signals in the high-gamma frequency range, which typically covers frequencies from 70 to 200 Hz.

A noticeable feature of acoustic speech signals is the fundamental frequency *f*_0_ of the human voice, which corresponds to the vibrational source of speech produced by the vocal folds in the larynx and further modulated by the vocal tract to produce the variety of speech sounds. The fundamental frequency depends on the size of the vocal folds and typically falls around 125 Hz for males and 215 Hz for women (Small, 2012). The high-gamma frequency band and the range of the fundamental speech frequency thus generally overlap.

Here, we analyzed human electrocorticographic (ECoG) and intracortical recordings during speech production and perception as well as rat microelectrocorticographic (µ-ECoG) recordings during sound perception. We found that electrophysiological signals are usually contaminated by spectrotemporal features of the sound produced by the participant’s voice or played by the loudspeaker. This contamination seems to result from a microphonic effect at the level of the cables and connectors along the recording chain, affecting the range of high-gamma frequencies and above. These findings suggest that care should be taken to avoid including these artifacts when investigating cortical signals underlying speech production and perception.

## 2 Methods

### 2.1 Human recordings

#### 2.1.1 Participants

The present study was conducted as part of the Brainspeak clinical trial (NCT02783391) approved by the French regulatory agency ANSM (DMDPT-TECH/MM/2015-A00108-41) and the local ethical committee (CPP-15-CHUG-12). It is based on electrophysiological recordings obtained in 3 patients at the Grenoble University Hospital: a 42-year-old (P2) and a 29-year-old (P3) males undergoing awake surgery for tumor resection, and a 38-year-old female (P5) implanted for 7 days as part of a presurgical evaluation of her intractable epilepsy. These 3 participants gave their informed consent to participate in the study. One additional 22-year-old male participant (HP) suffering from intractable epilepsy requiring surgical treatment has further been recorded at the second affiliated hospital of Zhejiang University. These procedures were followed from the guide and approved by the Second Affiliated Hospital of Zhejiang University, China. Participant HP gave written informed consent after detailed explanation of the potential risks of the research experiment.

#### 2.1.2 Electrophysiological recordings

Brain activity from participants P2 and P3 was recorded during awake surgery in the operating room just before tissue resection. For participant P2, a 256-electrode array (PMT Corp., USA) was positioned after opening the skull and the dura matter over the left sensorimotor cortex and the tumor (Figure 1, left). Ground and reference electrodes were integrated on the back side of the array and maintained wet using compresses soaked with saline. The 16 electrodes’ pigtails were connected to eight 32-channels Cabrio Connectors (Blackrock Microsystems, USA) connected by shielded cables to two front-end amplifiers (FEA, Blackrock Microsystems, USA) for amplification and digitalization at 10 kHz. The digitalized signals were then transmitted by an optic fiber to two synchronized Neural Signal Processors (NSP, Blackrock Microsystems, USA) interfaced with a computer. For participant P3, a 96-channel intracortical Utah microelectrode array (UEA, Blackrock Microsystems, USA) was inserted in the *pars triangularis* of Broca’s area (figure 1, middle), at a location that was subsequently resected to access the tumor for its removal. The pedestal serving as ground was screwed to the skull. Two wires with deinsulated tips were inserted below the dura, and one was used as reference. The electrodes were connected via a Patient Cable (Blackrock Microsystems, USA) to a FEA where signals were digitalized at 30 kHz and further transmitted through an optic fiber to a NSP.

**Figure 1.**
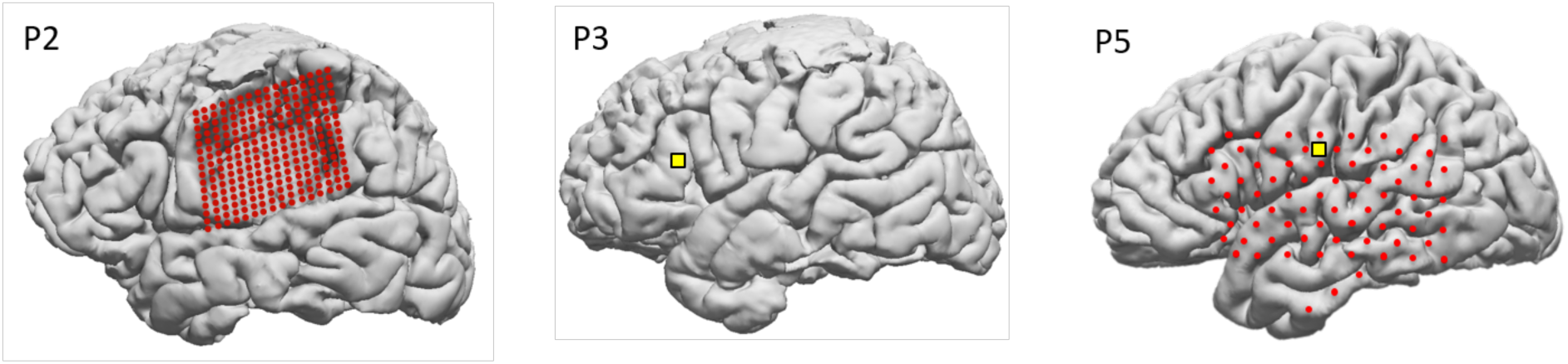
Electrode positions for human participants P2, P3 and P5. Red: ECoG electrodes (P2 and P5). Yellow: intracortical Utah array (P3 and P5).

Brain activity from participant P5 and HP was recorded in their room at the hospital. Participant P5 was implanted with a 72-electrode ECoG array (PMT Corp., USA) covering a large portion of her left hemisphere as well as a 4-electrode strip (PMT Corp., USA) over the left ventral temporal lobe and a 96-electrode UEA inserted in the left ventral sensorimotor cortex (Figure 1, right). Participant HP was implanted with a 32-electrode clinical subdural ECoG grid (HuakeHesheng, China) in his right sensorimotor cortex for clinical monitoring and localization of his seizure foci. The clinical electrodes were platinum electrodes with a diameter of 4 mm (2.3 mm exposed) spaced every 10 mm, implanted for seven days. The configuration and location of the electrodes, as well as the duration of the implantation, were determined by clinical requirements. In both participants, the transcutaneous pigtails of the ECoG grids were connected to PMT pigtail adaptors for P5 and HuakeHesheng adaptors for HP. These adaptors were connected to clinical headboxes (Blackrock Microsystems, USA) through individual touch-proof connectors and the headboxes to a FEA linked to a NSP. For participant P5, an electrode of the strip was used as the reference and another as the ground, and for participant HP, an electrode of the ECoG grid was used as the reference and another as the ground. For participant P5, the transcutaneous pedestal of the UEA was screwed to the skull and connected to a Cereplex-E headstage (Blackrock Microsystems, USA) ensuring signal amplification and digitization before transmission to a second NSP through a digital hub. For these intracortical recordings, the reference was a wire deinsulated at its tip and inserted below the dura, and the ground was the pedestal. For participant P5, data from both electrode arrays was sampled at 30 kHz and recorded on the two synchronized NSPs. For participant HP, the signals from the ECoG grid were sampled at 30 kHz.

#### 2.1.3 Audio recordings

For all participants, produced speech was recorded along with neural data. For participants P2, P3 and P5, a microphone (SHURE Beta 58 A) was positioned at about 10-20 cm from the mouth. The signal was amplified using an audio interface (Roland OCTA-CAPTURE) and digitized by one of the NSPs, at the same rate and synchronously with the neural data (see figure 2a). For participant HP, only the audio data of the sound file delivered to the patient was used in the present study.

**Figure 2.**
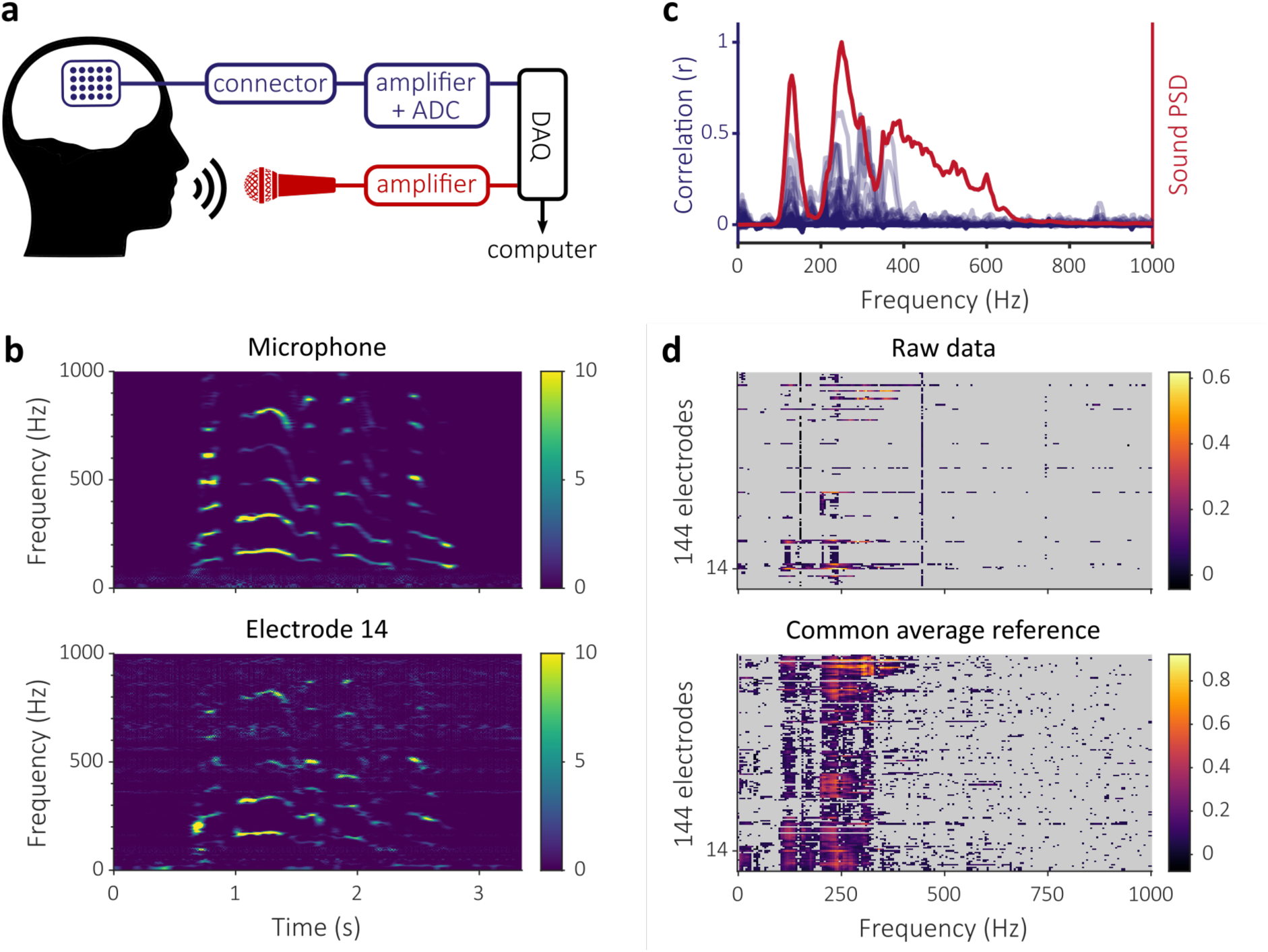
Correlation between voice and ECoG signals during speech production in participant P2. (a) Schematic representation of the recording setup, including neural (blue) and audio (red) data streams. The analog-to-digital conversion (ADC) of the audio signal is done in the data acquisition system (DAQ) whereas it in the FEA for neural signals (see section 2.1.2 for more details). (b) The upper and lower graphs show the z-scored spectrograms of the microphone and of electrode 14, respectively. The succession of stable striped patterns interleaved by transient states is typical of human speech formants. (c) Each blue curve represents, at all frequency bins, the value of the correlation coefficients between the spectrogram of one electrode signal and the spectrogram of the audio signal. The red curve represents the mean PSD of the audio signal (a.u.). (d) Heat maps representing the correlation coefficients between audio and neural data across electrodes and frequency bins. Correlation coefficients not statistically significant are displayed in grey. The upper and lower graphs show the results when using raw neural data and neural data after common average reference, respectively.

#### 2.1.4 Task and stimuli

All participants performed an overt speech production task. Particpants P2, P3 and P5 were asked to read aloud short French sentences, which were part of a large articulatory-acoustic corpus acquired previously (Bocquelet et al., 2016b) and made freely available (https://doi.org/10.5281/zenodo.154083). Participant P5 also took part in a protocol involving speech perception, where she was exposed to the sound of computer-generated vowels delivered by a loudspeaker positioned about 50 cm on her left. Participant HP was asked to listen and repeat aloud individual sentences of an ancient Chinese poem. Each block consisted of 4 sentences and each sentence lasted between 2 and 5 seconds. There were six blocks in total, 3 from the morning and 3 from the afternoon of the same day. The played sentences were realigned with the neural signals using triggers indicating the start of the stimuli.

### 2.2 Animal recordings

#### 2.2.1 Electrophysiological and audio recordings

In order to consider data recorded in a different condition, we also analyzed electrophysiological recordings obtained over the left auditory cortex of a ketamine (90 mg/kg)-xylazine (2 mg/kg) anesthetized 600-gram adult Sprague Dawley rat using a 64-electrode micro-ECoG array (E64-500-20-60-H64, NeuroNexus Inc, USA). These data were obtained in compliance with European (2010-63-EU) and French (decree 2013-118 of rural code articles R214-87 to R214-126) regulations on animal experiments, following the approval of the local Grenoble ethical committee ComEth C2EA-12 and the ministry authorization 04815-02. A bone screw was used for the ground and a stainless steel wire inserted below the skin ahead of Bregma was used for the reference. Signals were acquired using the RHD2000 acquisition system and two 32-channel RHD2132 headstages (Intan Technologies, USA). To avoid any possible crosstalk inside the Intan acquisition system, the sounds delivered to the rat were recorded on an independent CED Micro1401 (Cambridge Electronic Design, UK). Both acquisition devices were interfaced and synchronized by the Spike2 software with the IntanTalker module (CED programs) and signals were digitized at 33.3 kHz. The time jitter between sound and neural signals was checked to be below 2 ms.

#### 2.2.2 Audio stimuli

Pure tones (3-ms rise, 167-ms plateau and 30-ms fall times) with frequencies ranging from 0.5 to 16kHz were presented with pseudo-random inter-stimulus intervals of 1.8-2.2 seconds. Sounds were delivered at about 80-90 dB SPL in open field configuration using a MF1-S speaker (Tucker Davis Technology Inc, USA). The three lowest tone frequencies that were further considered in the present study are 0.5, 1 and 2.5 kHz.

### 2.3 *In vitro* recordings in PBS solution

A 24-electrode ECoG array (DIXI Medical SAS) was placed in a plastic container filled with a 1X phosphate-buffered saline (PBS). Two of the electrodes were used as the ground and reference electrodes, respectively. All electrodes were plugged into a clinical headbox (Blackrock Microsystems, USA) connected by shielded cables to a FEA and a NSP used for human recordings. A microphone was placed close to the plastic container. The audio data was acquired using the same hardware as for human P2-P3-P5 recordings (see section 2.1.3). Data was acquired at 30 kHz. Sounds were delivered either through a loudspeaker (M-Audio BX5-D2 loudspeaker used with participant P5) located about 1-2 m from the electrodes and recording chain, or very locally using a MF1-S speaker (Tucker Davis Technology Inc, USA) mounted in a closed-field configuration and sending sounds through a small plastic tube (3-mm outer diameter). A plastic box with a removable lid, soundproofed with cotton fiber insulation, was used to reduce sound propagation between the loudspeaker and the content of the box. For *in vitro* experiments, 20 pure tones lasting 4 seconds, with frequencies ranging from 25 to 975 Hz (spaced every 50 Hz), were played 4 times with 2 seconds of silence between sounds.

### 2.4 Data processing

#### 2.4.1 Data selection

10-min intervals with consistent speech production were selected from P2, P3 and P5 recordings. In the case of P5 recording in perception condition, the perception intervals had to be extracted, amounting to approximately 5.5 min. For animal recordings, a 10-min segment was selected. For each recording in PBS solution, the total duration of 9 minutes was kept for analysis. All recordings were visually inspected. For participant P2, 112 electrodes were removed due to several loose connections at the level of the Cabrio Connectors. For participant P5, 1 ECoG electrode showing saturating noise was removed. For participant HP, only the data during the speech perception stimuli were analyzed, which amounted for 68 seconds.

For all recordings, segments containing high-power transient noises were excluded from further analysis. To do so, the signal of each channel was detrended (using a 500-ms moving average) and positive and negative thresholds with an absolute value of 5 times the median absolute deviation were used to detect potential high-power noises. A sample was considered as noisy when at least 10% of the channels reached their threshold. A 500-ms window was also excluded around each noisy sample.

#### 2.4.2 Data pre-processing

A built-in analog band-pass filter was applied to the data recorded with the NSP (0.3-2500 Hz for 10 kHz sampling rate and 0.3-7500 Hz for 30 kHz sampling rate). Common average reference was applied only for comparison with original data as in figure 2d. To center audio signals, a moving average was computed over 1-second windows and subtracted.

#### 2.4.3 Spectrograms computation

In the present study, a spectrogram refers to the time-varying power spectral density (PSD) computed over a recording channel. For all analyses, spectrograms of neural and audio data were computed at a rate of 50 Hz using 200-ms time windows (weighted by a Hamming function). Mean sound PSDs were computed by averaging the spectrograms of audio signals over all selected time samples. For display purposes, the spectrograms in figure 2a and 2b were computed with higher frequency and time resolutions. These spectrograms were also z-scored within each frequency bin using artifact-free data segments containing the displayed extracts (60- and 30-second segments respectively).

#### 2.4.4 Spectrograms correlations

For all recordings, the correlations between the neural and the audio spectrograms were computed. For each electrode, the sample Pearson correlation coefficient *r* between the power amplitudes across time of the electrode and audio signals was computed for all frequency bins separately. For each value of *r*, a *p*-value was computed using Student’s *t*-test to test the null hypothesis that *r* = 0. The statistical significance of each correlation coefficient was then determined with respect to a Bonferroni adjusted significance level *α = 0.05/N* where *N* was the number of frequency bins times the number of electrodes in the recording.

### 2.5 Neural decoding

ECoG data from participants P2 and P5 were used to predict acoustic mel-cepstral coefficients of overt speech produced by the participants. Both participants were visually presented with a series of short sentences or vowel sequences written on a screen positioned about 50-100cm in front of them, and asked to repeat them overtly. The number of sentences was 118 for participant P2 and 150 for participant P5, corresponding to an overall duration of 230 and 329 sec of speech, respectively. The participants’ speech audio signals were decomposed into 25 mel-cepstral coefficients using the SPTK toolkit (*mcep* function). Spectrograms of the ECoG data were computed as described in paragraph 2.4.3 but at a rate of 100 Hz. Neural features were the spectrogram amplitudes in 10-Hz bands (i.e. 0-10, 10-20, … 190-200 Hz) and the band-pass filtered LFP signal (between 0.5 and 5 Hz). Two sets of neural features were considered, a first one where only features below 90 Hz were used and a second one where all features up to 200 Hz were used. A feature selection process was applied to keep only the features that were significantly modulated during speech production with respect to silence intervals, as assessed by Welch’s *t*-test with a Bonferroni adjusted significance level *α = 0.05/N* where *N* is the number of electrodes times the number of candidate features. The resulting number of selected features was 1413 out of 5376 for P2 and 648 out of 1512 for P5. These selected features were normalized and decomposed using PCA (both transformations were based on training sets). The first 50 (for the 0-90 Hz feature set) or 100 (for the 0-200 Hz feature set) components were used as the final set of features. A linear model was then used to map these neural features onto the mel-cepstral trajectories using 10-fold cross-validation. Chance decoding level was assessed by repeating the whole procedure after shuffling and time-reversing the mel-cepstral trajectories of the different sentences (truncation was applied to match sentence durations).

## 3 Results

### 3.1 Correlation between ECoG and sound signals during speech production

We observed strong correlations between ECoG and sound spectrograms in participant P2 during speech production. Participant P2’s brain activity was recorded with an ECoG grid while he was reading sentences aloud. Simultaneously, a microphone was used to capture the sound of his voice (see figure 2a). Figure 2b shows a portion of the z-scored spectrograms of the sound signal (top) and of an electrode of the ECoG grid (bottom). In this example, the ECoG signal shows a very similar spectrotemporal structure as that of the sound. The time-frequency patterns observed are consistent with human speech and are unlikely to be brain activity. This is actually further assessed by *in vitro* tests described below (see section 3.4), which show such strong correlations when electrodes are simply immersed in PBS. We therefore attribute the high degree of similarity between the two bio-signals to a contamination of the electrophysiological measure.

We quantitatively assessed this phenomenon by computing the correlation between the power of the signal within each frequency bin of each electrode signal with that of the sound signal. As shown in figure 2c and in the top of figure 2d, correlations up to 0.6 could be observed depending on the electrode. Up to 370 Hz, the strongest correlations were observed at frequencies most present in the sound signal, and in particular between 115 and 145 Hz, which corresponded to the range of the fundamental frequency of the subject’s voice. This correspondence between the peaks of the mean sound PSD and the frequency bins showing high correlations supports the hypothesis that at least part of the correlation is caused by acoustic contamination of the neural data. Above 370 Hz, correlations were low even at frequencies for which the power of the speech signal remained high. As shown in figure 2d, the correlations between sound and ECoG spectrograms were still present and even exacerbated after common average re-referencing of the ECoG signals.

We carried out the same analysis for the ECoG data of participant P5 during speech production (figure 4a, top). In this case, the significant correlation coefficients reached values up to 0.2. Most of the high-valued coefficients were observed in a narrow band around 225 Hz, which corresponded to the range of the fundamental frequency of the participant’s voice.

### 3.2 Correlation between intracortical and sound signals during speech production

In P3 recording, we further observed statistically significant correlations between the spectrograms of intracortical signals recorded using a Utah array and that of the produced speech signal. Figure 3a shows a portion of the z-scored spectrograms of the subject’s voice (top) and of one electrode of the array (bottom). The spectrogram of the selected micro-electrode clearly shows spatio-temporal features also observed in the sound spectrogram (between 200 Hz and 400 Hz). Statistically significant correlation coefficients up to 0.7 were observed, with peaks falling in the range of frequencies where the sound signal showed high power (figure 3b). Noticeably, correlations between intracortical and sound signals during speech production were much weaker in participant P5 (figure 4b).

**Figure 3.**
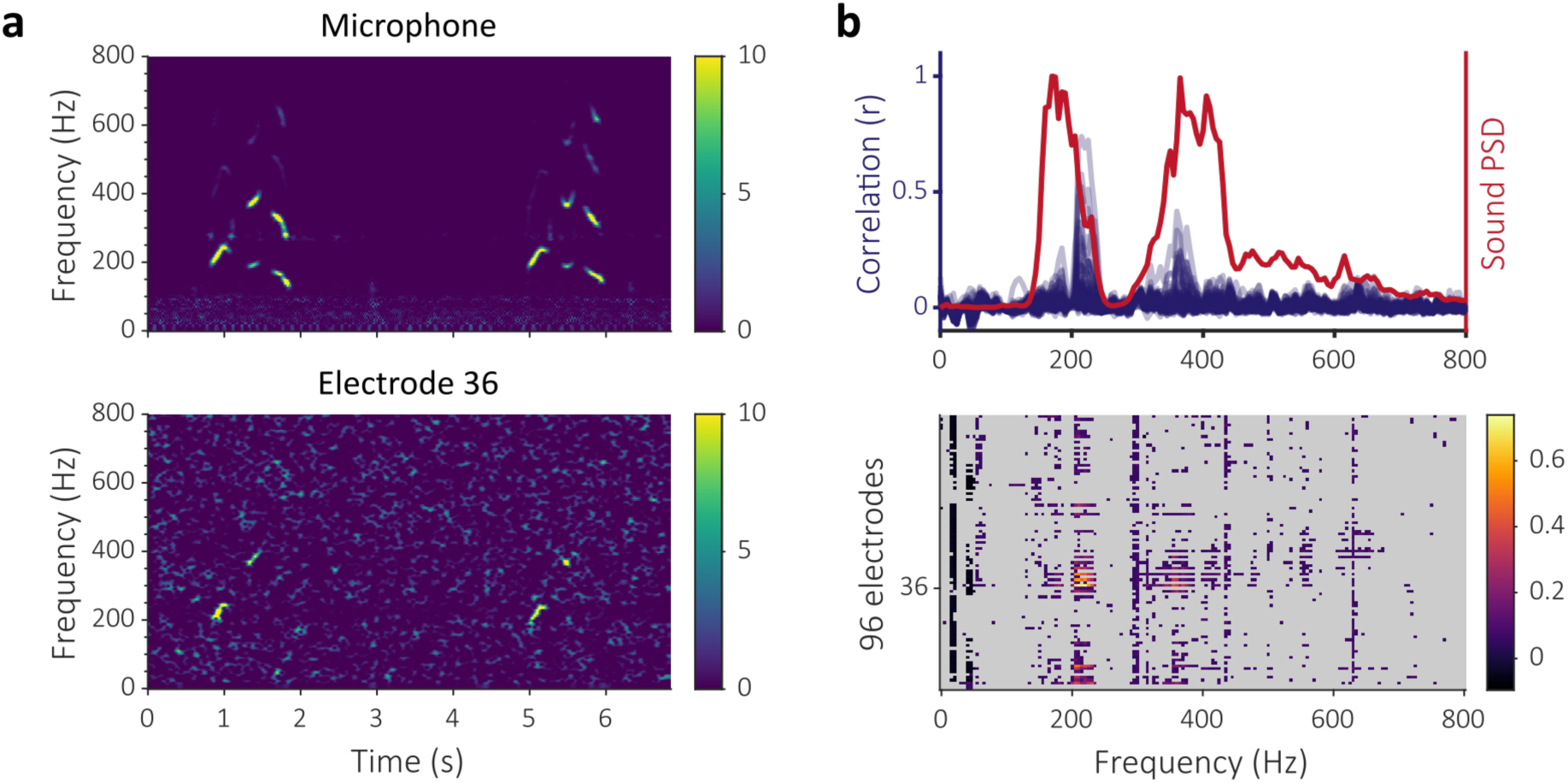
Correlations between voice and intracortical signals during speech production in participant P3. (a) The upper and lower graphs show the z-scored spectrograms of the microphone and electrode 36, respectively. (b) On the upper graph, each blue curve represents, at all frequency bins, the value of the correlation coefficients between the spectrogram of one electrode signal and the spectrogram of the audio signal. The red curve represents the mean PSD of the audio signal (a.u.). The lower panel represents a heat map of the correlation coefficient between audio and neural data for all electrodes and frequency bins. Correlation coefficients not statistically significant are displayed in grey.

**Figure 4.**
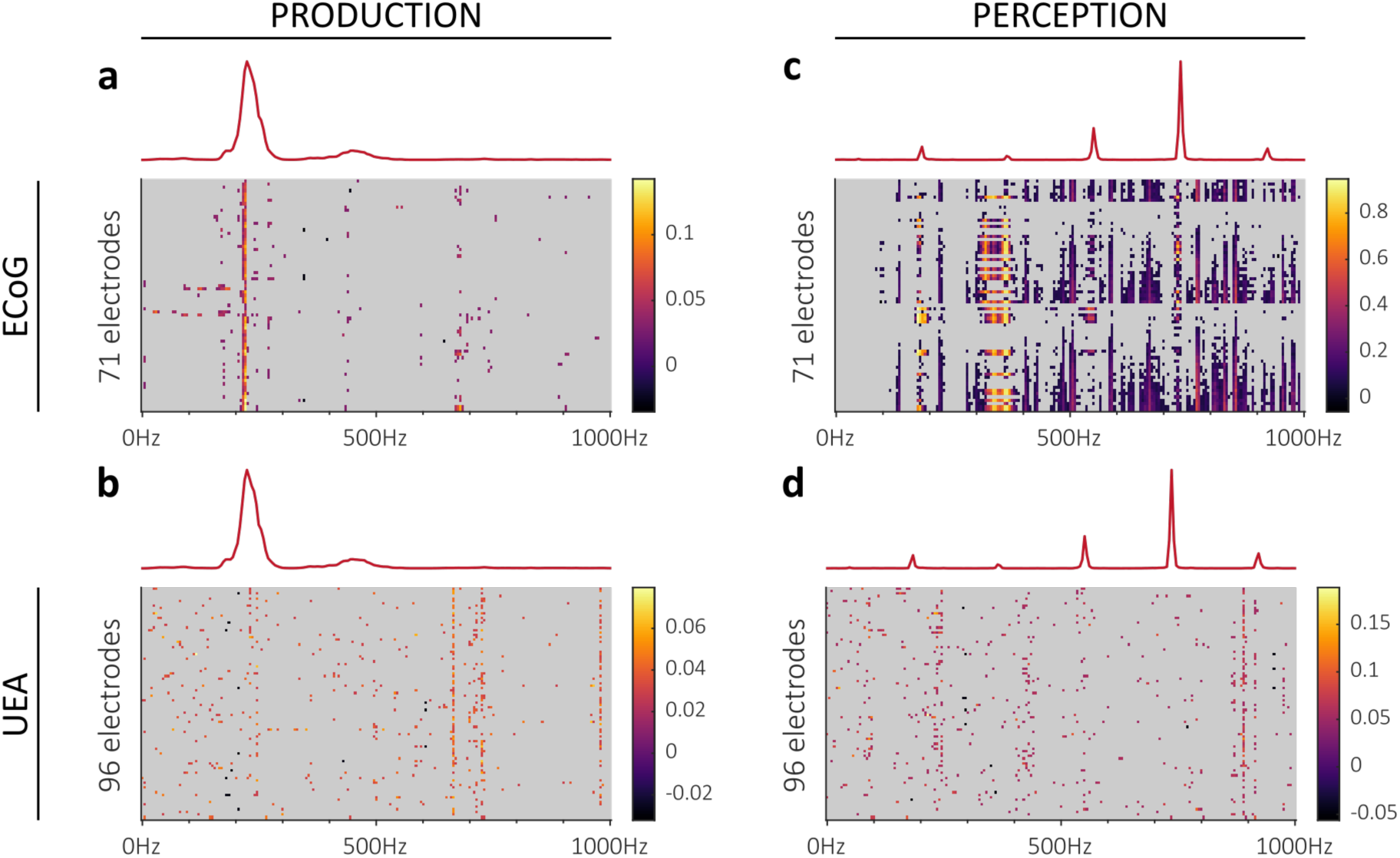
Spectrogram correlations between sound and neural data observed in participant P5. The results are presented depending on the experimental condition (speech production or perception) and the type of electrophysiological measure (ECoG and UEA). The heat maps show the value of the correlation coefficient with the spectrogram of the sound for each electrode and each frequency bin. The correlation coefficients that are not statistically significant are displayed in grey. The red curves indicate the mean PSD of the sound (a.u.) recorded during the experiments. The speech perception experiment used computer-generated vowels, peaks appearing in the mean PSD are the fundamental frequency and first harmonics of the vocal synthesizer. (a, b) Results for the speech production condition using ECoG and UEA data, respectively. (c, d) Results for the speech perception condition using ECoG and UEA data, respectively.

### 3.3 Correlation between electrode and sound signals during sound perception

Statistically significant correlations between electrode and sound signals were not only present during speech production as reported above, but also during sound perception. This phenomenon was observed in human and animal recordings using completely different recording instrumentations.

#### 3.3.1 Human recordings

Participant P5 also participated in a paradigm where artificially synthesized speech sounds were presented to her through a loudspeaker positioned on her left. Brain activity was recorded from both ECoG electrodes and intracortical microelectrodes. The sound produced by the loudspeaker was also recorded simultaneously. Performing the same analysis as for speech production data, we found that ECoG signals showed strong correlations with the sound signal, with peaks up to 0.9 (figure 4c). As observed in recordings during speech production, frequencies showing strong correlations are mostly found in the bands that concentrate most of the sound power. These bands correspond mainly to the pitch of the synthesized sound (185 Hz) and its harmonics. By comparison, the spectrograms of intracortical signals were poorly correlated with that of the sound (figure 4d).

#### 3.3.2 Rat recordings

In order to verify that the correlations were not due to our clinical recording system in particular, we performed the same type of analysis on data obtained from an experiment in a rat. The left auditory cortex was recorded using a commercial µ-ECoG grid connected to an Intan neural recording system (figure 5a). In this case, pure tones were delivered in an open field configuration. As shown in figure 5b, we again observed strong correlations between the electrode and sound spectrograms, with sharp peaks at the specific frequencies of the pure sound stimuli (500, 1000 and 2000 Hz).

**Figure 5.**
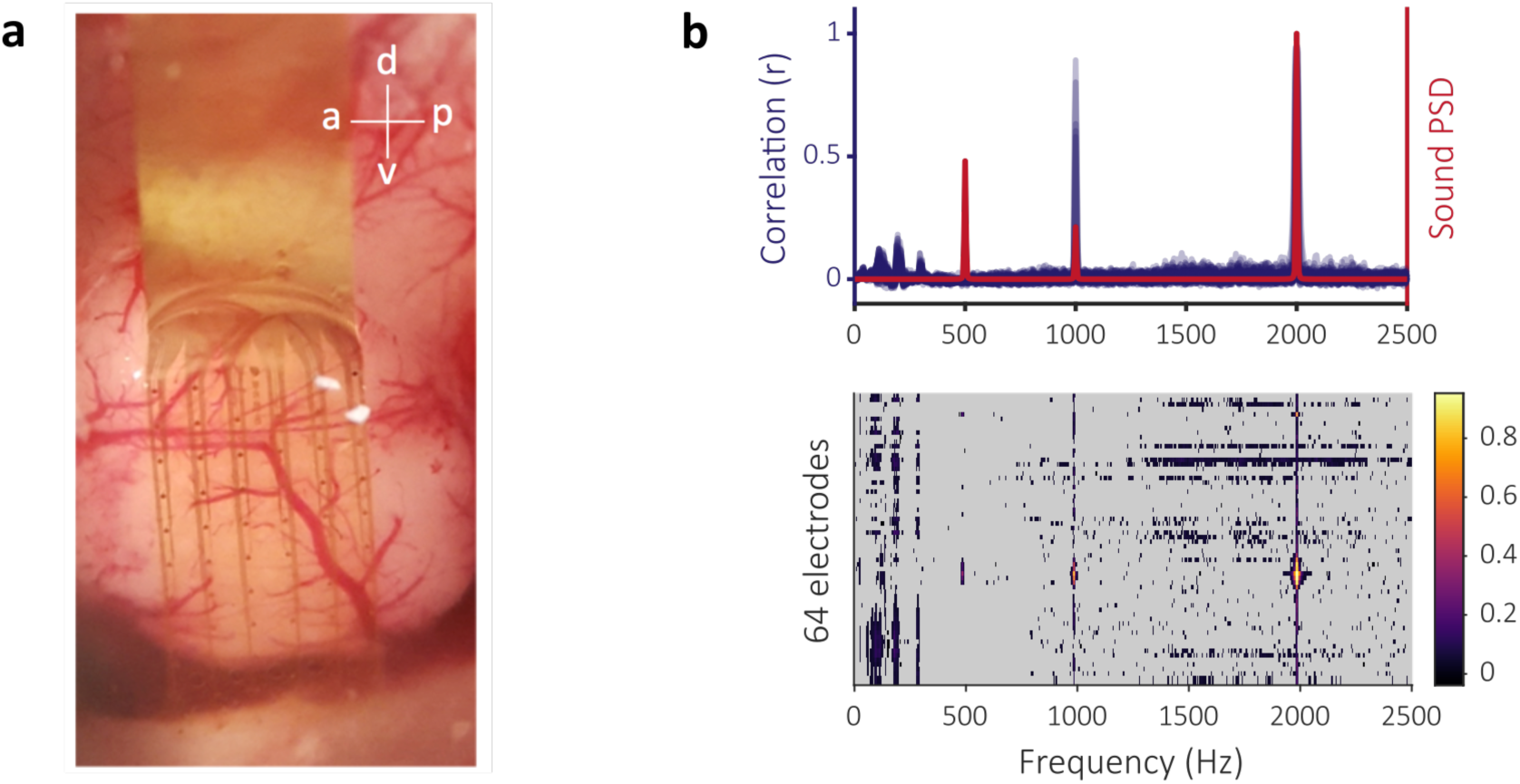
Correlations between sound and µ-ECoG spectrograms during pure tones perception in an anesthetized rat. (a) Photograph of a µ-ECoG grid positioned over the left auditory cortex of a rat. Directions: a = anterior, p = posterior, d = dorsal, v = ventral. (b) On the upper panel, each blue curve represents, at all frequency bins, the value of the correlation coefficients between the spectrogram of one electrode signal and the spectrogram of the audio signal. The red curve represents the mean PSD of the audio signal (a.u.). The lower panel represents a heat map of the correlation coefficient between audio and neural data for all electrodes and frequency bins. Correlation coefficients not statistically significant are displayed in grey.

### 3.4 Potential influence of contamination on speech decoding

We assessed the potential influence of contamination of electrophysiological signals by sound on the performance of neural decoding to predict acoustic features of produced speech. For this purpose, we considered ECoG data from participant P2 which shows strong evidence of contamination (Figure 2b), and ECoG data from participant P5 which showed weak or no contamination below 200 Hz (Figure 4). As illustrated in Figure 6, we found that including features up to 200 Hz resulted in a drastic increase in decoding performance for P2 (compare the green plots of Figures 6a versus 6b), and only limited improvement for participant P5 (Figures 6c versus 6d).

**Figure 6.**
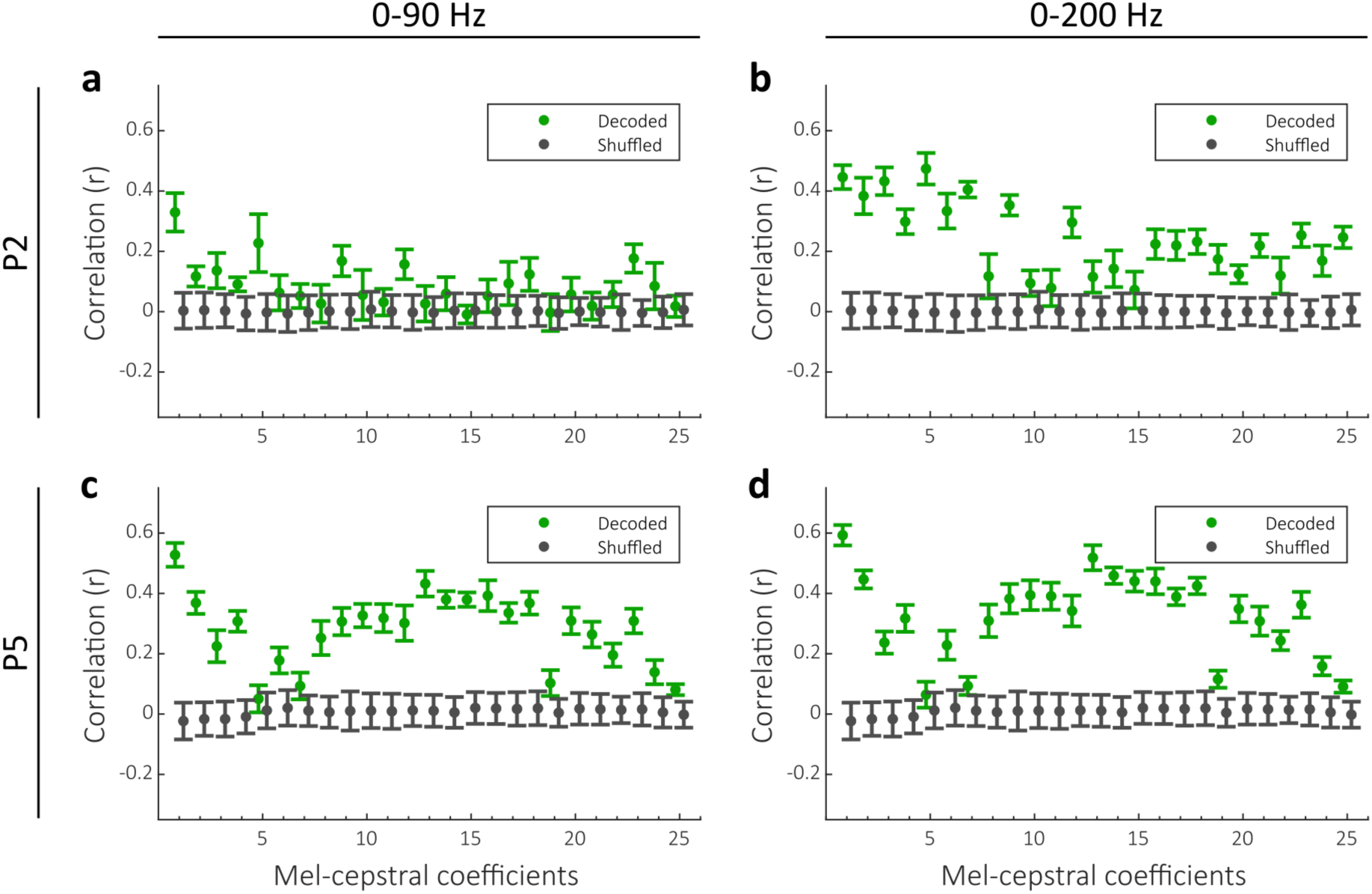
Linear decoding of acoustic coefficients of overt speech for participant P2 (panels a and b) and P5 (panels c and d). The ECoG spectral features ranged either between 0 and 90 Hz (a,c) or between 0 and 200 Hz (b,d).

### 3.5 Sound contamination and electrode quality

In the following of this paper, we investigate the possible causes of the contamination observed in neural signals. We first tested whether the level of sound contamination was determined by the quality of the electrode signal. Participant HP was recorded twice on a single day, once in the morning and once in the afternoon. Between the two sessions, the electrodes were disconnected and then reconnected after lunch. Sound contamination was observed only in the afternoon session (Figure 7a-b), which indicates that the contamination was not related in this case to the electrode array and its intracranial environment as those remained unchanged. Moreover, electrodes showing strong contamination in the afternoon session showed very variable signal quality (Figure 7c). For instance, one electrode with very strong 50-Hz power-line noise (considered as a typical “bad channel”) showed a strong contamination, while two other channels with no such noise showed an equally strong contamination for one, and very weak or no contamination for the other. These observations indicate that the quality of the signal was not a sufficient predictor of sound contamination.

**Figure 7.**
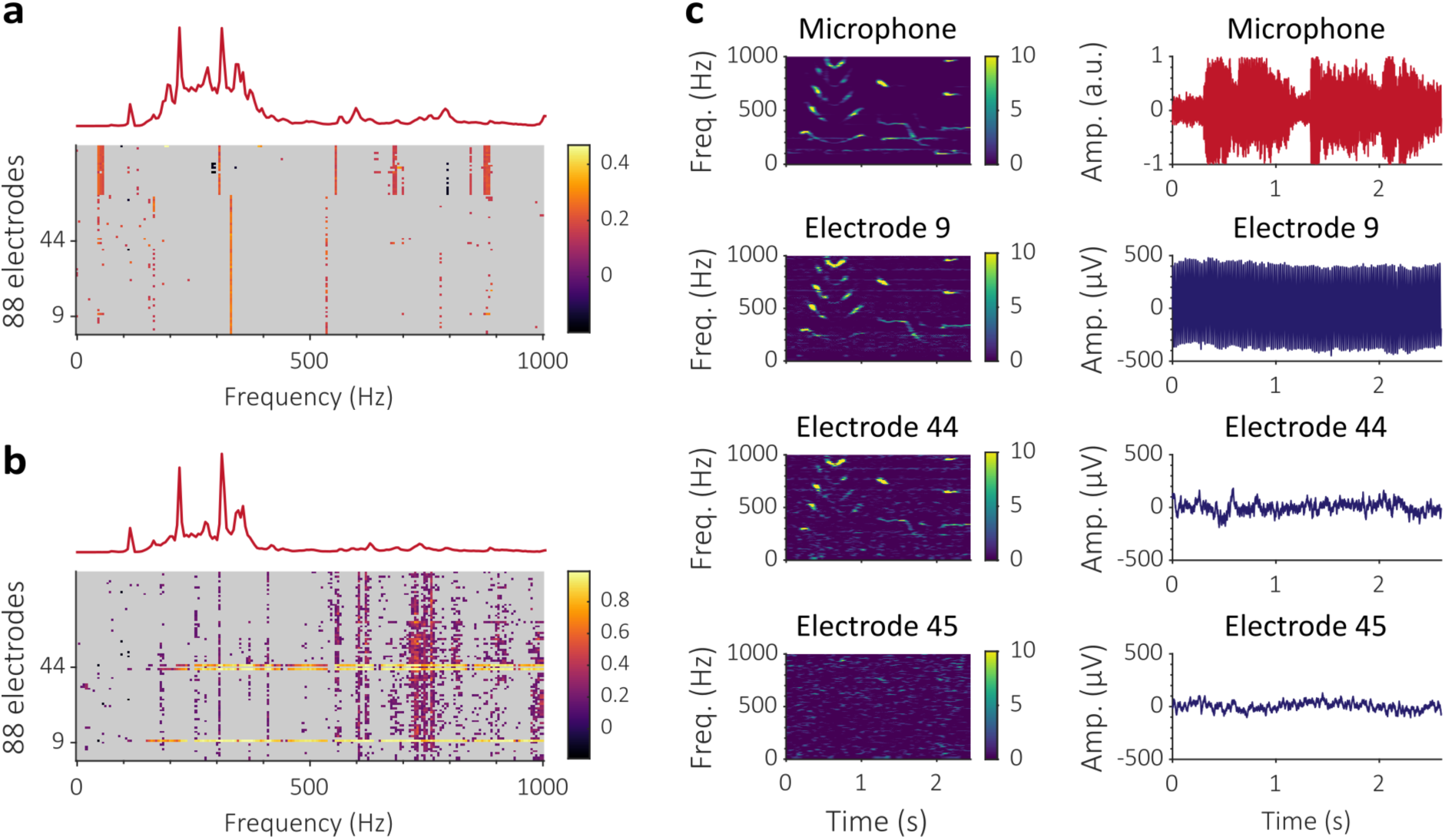
Correlations between sound and ECoG recordings in participant HG during speech perception. Correlation heat maps (as in Figure 4) for the morning (a) and afternoon (b) sessions. (c) Example of 2.5-sec spectrograms (left) and raw signals (right) for the sound that was presented to the participant and signals from 3 electrodes of the grid, one with high 50-Hz power-line noise (electrode 9), and two with standard noise level (electrodes 44 and 45).

### 3.6 *In vitro* evidence of acoustic contamination

Next, we used a reduced experimental setup to determine more in details the cause of the observed correlations (see figure 8). The experiment was designed to verify that the correlations between the sound and the electrode recordings can be obtained without brain activity and to attempt to demonstrate that the correlations originate from the mechanical transmission of sound vibrations. The electrical potentials of ECoG electrodes placed in PBS were recorded while pure tone sounds were played by a loudspeaker. In order to evaluate the intensity of the incident sound, a microphone was placed near the container filled with PBS. A soundproof box was used to insulate either the loudspeaker, the ECoG array, or part of the acquisition chain. The function of the box was to reduce the propagation of sound from the loudspeaker to the devices without substantially interfering with other parameters of the experiment. To determine the impact of sound propagation on the spectrogram correlations, we analyzed the data in open and closed box conditions.

**Figure 8.**
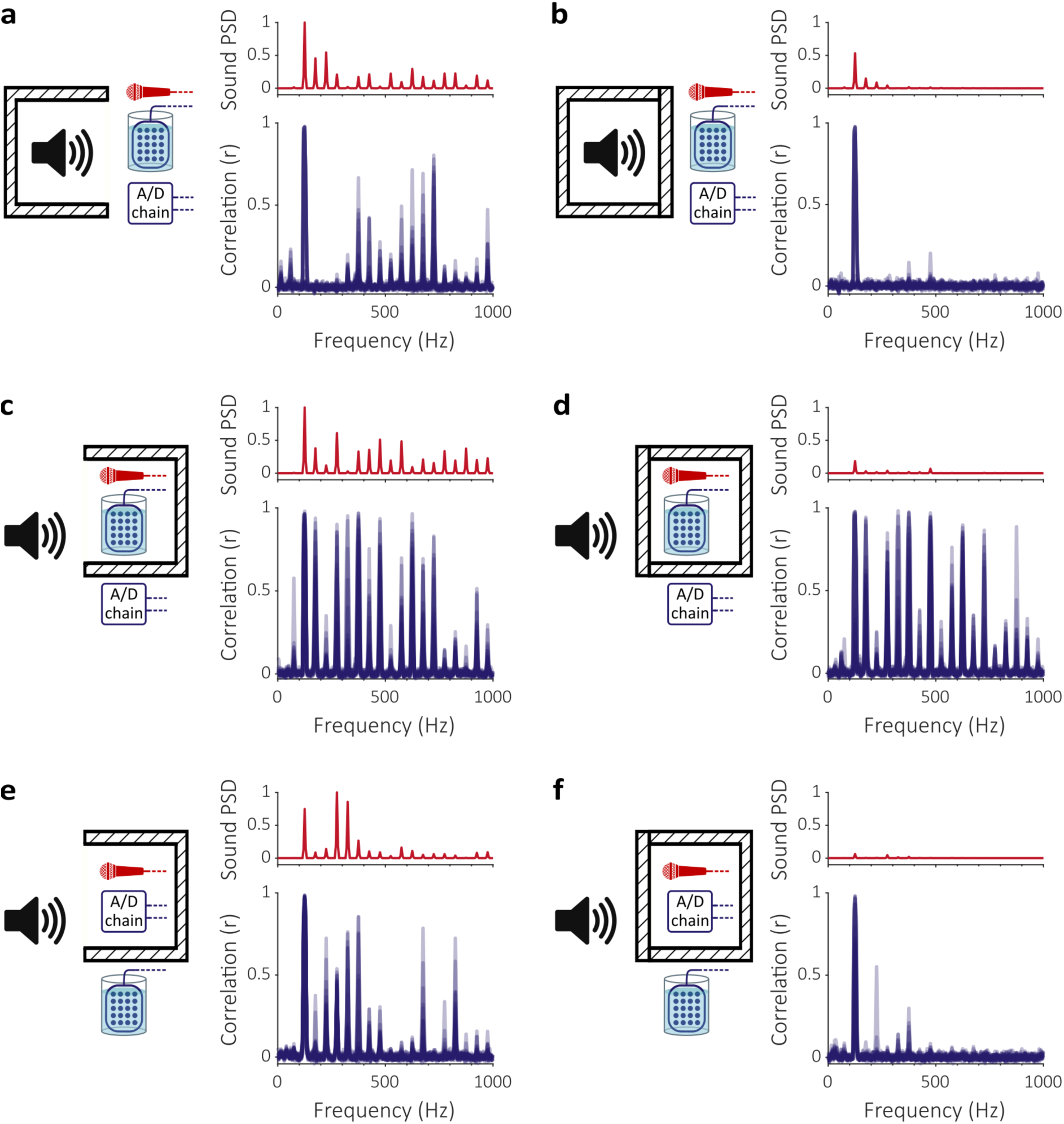
Correlations between sound and ECoG recordings in different in vitro experimental configurations. The red graphs show the mean PSD of the sound captured by the microphone. Mean PSD scale (a.u.) is common within each of the 3 panel pairs a-b), c-d) and e-f). In the blue graphs, each curve represents, at all frequency bins, the value of the correlation coefficients between the spectrogram of one electrode signal and the spectrogram of the audio signal. (a) Loudspeaker placed in the open box. (b) Loudspeaker placed in the closed box. (c) Electrodes and microphone placed in the open box. (d) Electrodes and microphone placed in the closed box. (e) Amplification and digitization chain (composed of the cables, adaptors, splitter box and FEA) and microphone placed in the open box. (f) Amplification and digitization chain and microphone placed in the closed box. Within each raw, only the fact that the lid was open or closed changed.

In the first configuration, the loudspeaker was placed in the open box (figure 8a). As for *in vivo* experiments, we found that high correlations occurred at some of the frequencies of the sound stimuli. For some electrodes, the value of the correlation coefficient at 125 Hz was larger than 0.9. This result demonstrates that spectrogram correlations similar to those described in sections 3.1–3.3 occur in absence of any brain activity. In the second configuration, the loudspeaker was placed in the closed box (figure 8b). The reduction of the power of the incident sound due to the insulation is confirmed by the mean sound PSD (figure 8b, top). We observed that most of the correlation coefficients also have much lower values (figure 8b, bottom – compare with figure 8a, bottom). This result supports the hypothesis of acoustic contamination, i.e. that the spectrogram correlations between sound and electrodes data originate from the mechanical propagation of sound to the neural recording hardware.

In the third and fourth configurations, the electrode array and the microphone were placed in the box but the rest of the acquisition chain was left outside. When the box was left opened (figure 8c), we observed high correlations at the frequency of the stimuli, similarly to the previous open box condition (figure 8a). The differences of frequency responses visible in the mean sound PSD across the 3 open box conditions can be explained by the modification of the arrangement of the experimental setup. In the last configuration, the box was closed over the electrodes and microphone (figure 8d). The sound insulation provided by the box was confirmed by the large reduction of the sound stimuli mean PSD (figure 8d, top). However, as shown in the bottom graph of figure 8d, the spectrogram correlations remained largely unaffected by the closing of the lid over the electrode array, contrarily to the previous experiment where the lid was closed over the loudspeaker (figure 8b). This suggests that the acoustic contamination of the electrical potential measurement may not only occur at the electrode level but also at other levels of the acquisition chain.

To test this hypothesis, the amplification and digitization chain (A/D chain, composed by the cables, adaptors, splitter box and FEA) was put inside the sound-attenuating box with the microphone. In this case the electrodes in PBS were outside the box. While correlations were high when the box was open (figure 8e), they were strongly reduced when the box was closed (figure 8f). This further confirmed that the acoustic contamination mainly occurs in the recording chain and not at the electrodes level.

### 3.7 Localization of acoustic contamination along the recording chain

Finally, we aimed at determining where along the recording chain the contamination occurred. The fact that contamination was observed in participant HP in the afternoon but not in the morning session suggests that disconnecting and reconnecting the electrodes to the system could have produced the contamination to occur in the afternoon. To test this more thoroughly, sounds were delivered very locally at different locations along the recording chain connected to electrodes bathed in PBS (Figure 9). No contamination could be observed when the sound was delivered next to the electrodes in the PBS solution. By contrast, a contamination was observed when the sound was delivered against the ECoG grid cables, the pigtail connectors, or the headbox touchproof connectors. No contamination occurred along the shielded ribbon cable downstream of the headbox or at the level of the FEA.

**Figure 9.**
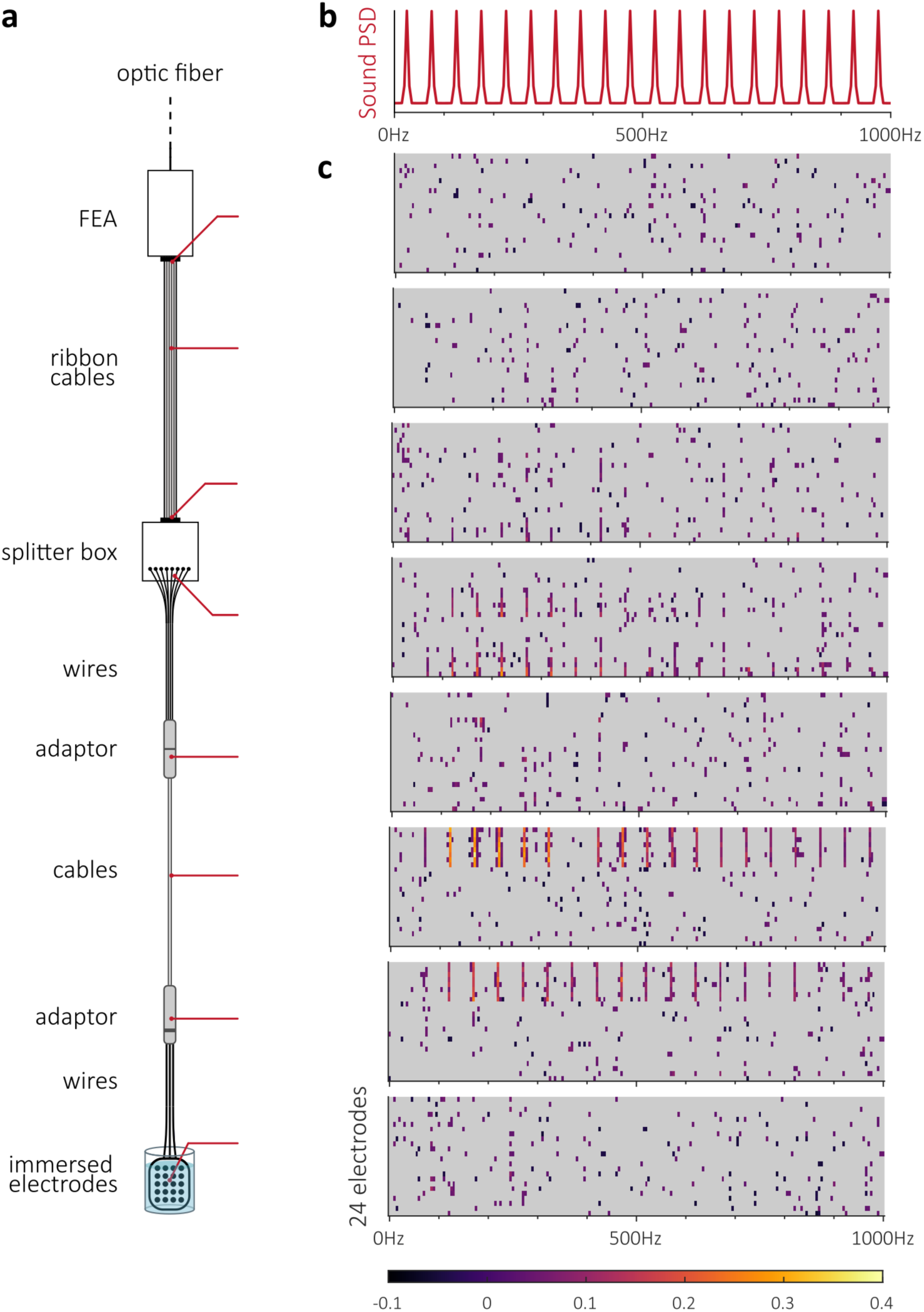
Determination of the location of sound contamination along the recording chain (case of ECoG electrodes). a) Sounds were delivered focally at different locations (indicated by the red lines) of the recording chain. b) PSD of the sounds delivered through the speaker. c) Correlation heat maps for each location of sound delivery (each map is displayed against the corresponding location of sound delivery indicated in panel a).

## 4 Discussion

Data considered in this study includes human and animal recordings during speech production and/or sound perception tasks. Using these different setup conditions, we observed statistically significant correlations between spectrograms of electrophysiological and simultaneously recorded audio signals. These correlations occurred at frequencies most present in the sound signal. The recordings used ECoG and intracortical micro-electrode arrays, interfaced with different data acquisition systems. Based on the present findings, it is not possible to draw quantitative conclusions about the influence of these factors, but the variety of cases suggests that the presence of the observed correlations is a widespread phenomenon.

Motion artifacts are classically seen in electrophysiological signals. In particular, mechanical vibrations may create variations of biopotential measurements (Luna-Lozano and Pallas-Areny, 2010). Such undesired signals may have different origins, including the bending of the electrode wires and the electrochemical changes at the electrode-electrolyte interface induced by small displacements of the electrodes (Salatino et al., 2017; Nicolai et al., 2018). Here we observed sound contamination of neural signals in different setups. We could reproduce the phenomenon in a minimal *in vitro* setup, confirming that sound-electrode correlations do not originate from brain activity and arise from the impact of sound vibrations on the acquisition chain. The experiments shown in section 3.4 further suggest that in the tested setup, the microphonic effect does not necessarily take place at the electrodes’ level, but in the rest of the recording chain. Focal sound delivery at different locations along the chain allowed to precisely identify that acoustic vibrations of the sound itself contaminated the electrode signals at the level of unshielded cables and connectors. Improving these parts of the recording chain thus appears mandatory to ensure proper and artifact-free neural signal recordings.

The extent to which the acoustic noise spectrally overlaps with the measured brain activity depends on the nature of the sound and on the studied activity. In the case of ECoG recordings during speech production paradigms (see section 3.1), the overlap between the range of the voice fundamental frequency and the high-gamma band might make it difficult to record an artifact-free signal in this band. As suggested by results in section 3.3, sound stimuli, and by extension any sound during the recording, could contaminate the recorded data in any frequency band. It can thus be expected that high-frequency components of the sound might also influence the detection of multi-unit activity in micro-electrode recordings (see section 3.2).

In conclusion, the purpose of this study is to alert on possible microphonic contamination of neural signals so that care could be taken to evaluate and eliminate this problem, especially when building decoders of neural activity underlying overt speech production or sound perception. Experimental setups might be improved to become less sensitive to microphonic effects, and signal-processing techniques might be developed to eliminate sound contamination in neural recordings. It is very important to underline that this report does not question the existence of relevant physiological neural information in high-gamma frequency signals underlying speech production or sound perception. Indeed, it has been shown by several groups that spectral features of imagined speech or silent articulation can be predicted from low or high-gamma signals recorded in participants not overtly speaking (Pei et al., 2011; Ikeda et al., 2014; Martin et al., 2014, 2016; Bocquelet et al., 2016a; Anumanchipalli et al., 2019; Gehrig et al., 2019). Future developments of speech prostheses should thus build upon these findings.

## Acknowledgments

We thank C. Kell (Universitätsklinikum Frankfurt), and P. Megevand (Univ Geneva) for fruitful discussions and for making available additional ECoG data not presented in this manuscript on which similar analyses could be applied. This work was supported by the FRM Foundation under grant No DBS20140930785, the French National Research Agency under Grant Agreement No. ANR-16-CE19-0005-01 (Brainspeak), and by the European Union’s Horizon 2020 research and innovation program under Grant Agreements No. 732032 (BrainCom), No. 696656 (GrapheneCore1), and No. 785219 (GrapheneCore2).

